# *De novo* basecalling of m^6^A modifications at single molecule and single nucleotide resolution

**DOI:** 10.1101/2023.11.13.566801

**Authors:** Sonia Cruciani, Anna Delgado-Tejedor, Leszek P. Pryszcz, Rebeca Medina, Laia Llovera, Eva Maria Novoa

## Abstract

RNA modifications hold pivotal roles in shaping the fate and function of RNA molecules. Although nanopore sequencing technologies have proven successful at transcriptome-wide detection of RNA modifications, current algorithms are limited to predicting modifications at a per-site level rather than within individual RNA molecules. Herein, we introduce *m^6^ABasecaller*, an innovative method enabling direct basecalling of m^6^A modifications from raw nanopore signals within individual RNA molecules. This approach facilitates *de novo* prediction of m^6^A modifications with precision down to the single nucleotide and single molecule levels, without the need of paired knockout or control conditions. Using the *m^6^ABasecaller*, we find that the median transcriptome-wide m^6^A modification stoichiometry is ∼10-15% in human, mouse and zebrafish. Furthermore, we show that m^6^A modifications affect polyA tail lengths, exhibit a propensity for co-occurrence within the same RNA molecules, and show relatively consistent stoichiometry levels across isoforms. We further validate the *m^6^ABasecaller* by treating mESC with increasing concentrations of STM2457, a METTL3 inhibitor as well as in inducible METTL3 knockout systems. Overall, this work demonstrates the feasibility *de novo* basecalling of m^6^A modifications, opening novel avenues for the application of nanopore sequencing to samples with limited RNA availability and for which control knockout conditions are unavailable, such as patient-derived samples.

## INTRODUCTION

RNA modifications, also known as epitranscriptomic modifications, are chemical alterations that occur on RNA molecules, affecting their fate and function. They can play crucial roles in a wide range of biological processes, such as cellular differentiation ^1–3^, immune response ^4,5^, cancer progression ^6,7^ and sex determination ^8,9^. At the molecular level, they can alter gene expression ^10^, protein translation ^11^, RNA stability ^12^ and RNA-protein interactions ^13^.

More than 170 different RNA modifications have been described to date ^14^. All four RNA bases, as well as the sugar moiety, can be targets of modification, and all known RNA species are subject to modifications, including ribosomal RNAs (rRNAs), messenger RNAs (mRNAs), transfer RNAs (tRNAs) and small non-coding RNAs (snRNAs). Most attention has been focused on characterising N6-methyladenosine (m^6^A), the most abundant internal modification in eukaryotic mRNA molecules ^15,16^, which can be dynamically regulated upon a broad range of conditions and environmental stimuli ^17^. m^6^A modifications are deposited by “writers” that catalyse the addition of the methyl group ^18,19^, they can also be removed by “erasers” of the modification ^20,21^, and recognized by “readers” that will selectively bind to the modified RNA ^22–25^. The study of the function and dynamics of m^6^A modifications has been greatly facilitated by the use of next-generation sequencing (NGS) technologies, which have enabled transcriptome-wide mapping of m^6^A sites across species, cell types and environmental conditions ^26–32^.

Nanopore direct RNA sequencing (DRS) has recently emerged as a promising alternative to NGS-based methods to comprehensively investigate the epitranscriptome ^33–36^. This technology capitalizes on the measurement of current intensity fluctuations generated as the RNA or DNA molecules translocate through the nanopores. In contrast to NGS approaches, nanopore DRS can generate transcriptome-wide maps of RNA modifications at the isoform level ^37^, can simultaneously detect diverse RNA modification types ^38,39^ and provide quantitative estimates of RNA modification levels at individual sites ^40,41^. Consequently, it empowers the generation of transcriptome-wide epitranscriptomic maps at an unprecedented resolution–unattainable through short-read sequencing data.

In recent years, several studies have showcased the capacity of nanopore DRS to detect diverse RNA modification types ^42,43^, encompassing m^6^A ^44–51^, pseudouridine ^38,52,53^ and inosine ^54^. To identify RNA modifications in nanopore DRS data, two major strategies have been employed: (i) identifying modifications through scrutinising current intensities and/or dwell times ^38,49,51^; and (ii) identifying modifications through the analysis of basecalling ‘errors’ ^44,55–58^ (depicted in **Figure 1A**). While both approaches yield success at detecting RNA modifications, they typically require comparing the reads of interest with those from a reference sample, such as a knockout or knockdown condition of the RNA modification ‘writer’ enzyme of interest. This need of ‘paired’ conditions greatly limits the applicability of nanopore DRS, as every study must first obtain a knockout or knockdown condition of the ‘writer’ enzyme in their model system prior to sequencing their samples. To overcome this limitation, methods that circumvent the need for ‘paired’ conditions have been recently developed ^49^; however, the predictions of these methods are not at per-molecule level, but rather, at per-site resolution.

**Figure 1.**
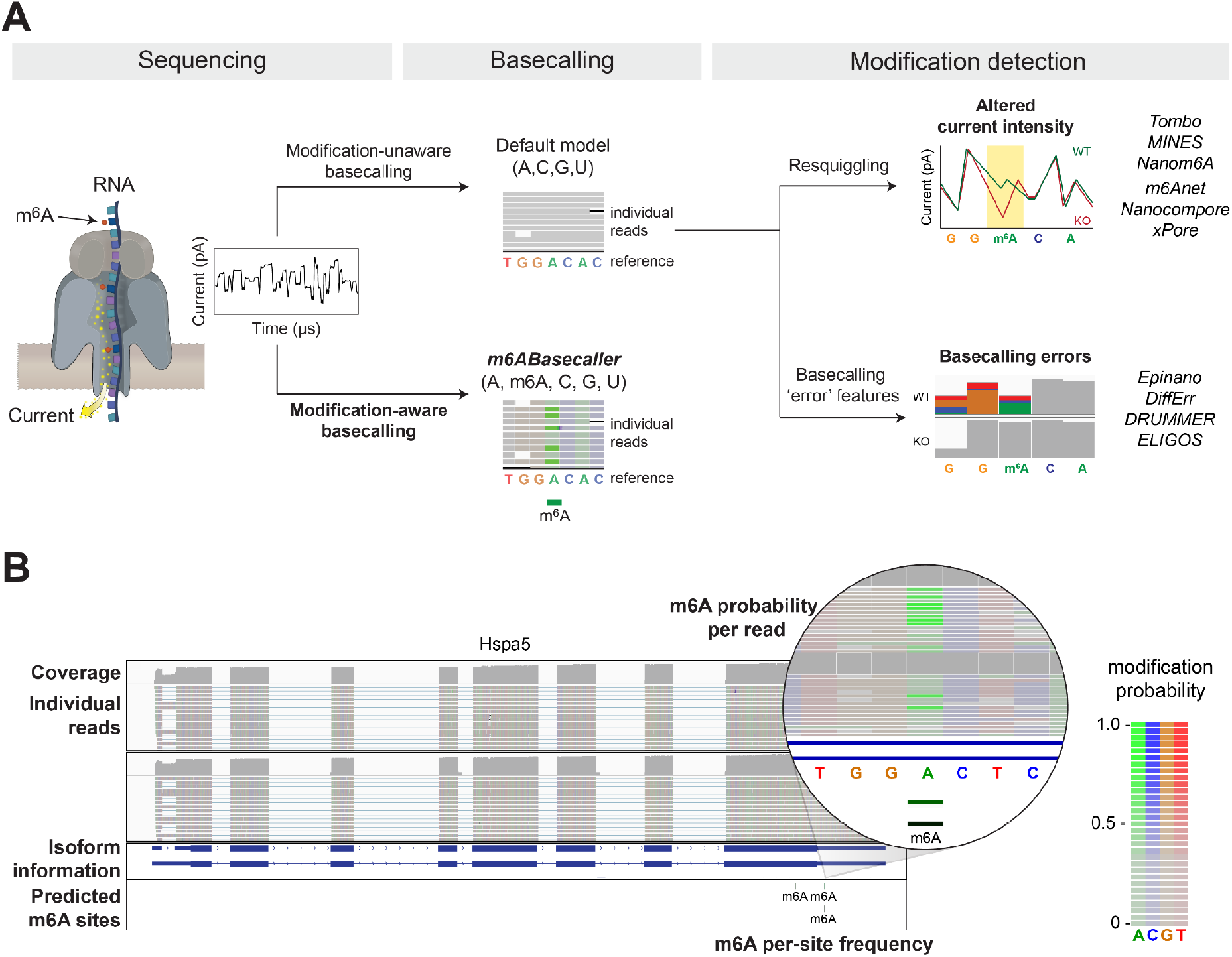
Schematic overview of the approaches that can be used to identify RNA modifications from direct RNA sequencing (DRS) data. **(A)** Overview of the methods used to detect modified sites from DRS data. Commonly used softwares to detect RNA modifications rely on either: i) basecalling errors that are present in a wild type (WT) but not a knockout (KO)/control condition, or ii) altered current intensities when comparing WT and KO/control conditions. All these methods use default (modification-unaware) RNA basecalling models and require extensive post-processing after basecalling steps –mapping, resquiggling, feature extraction and statistical testing– to identify modified sites. The alternative option is to use a modification-aware RNA basecalling model that predicts modifications during the basecalling step, which provides m^6^A modification predictions with single nucleotide and single molecule resolution. **(B)** IGV visualisation of a BAM file where reads have been basecalled using the *m^6^ABasecaller*, allowing per-read analysis of m^6^A modifications in full-length reads. BAM files have m^6^A information encoded at per-read and per-nucleotide level in the form of modification probabilities. Colouring nucleotides based on their modification probability allows simple visualisation of m^6^A-modified sites (bright green) in a transcriptome-wide fashion. A ‘predicted m^6^A site’ is defined as a position that has at least 25 reads coverage and ≥ 5% modification stoichiometry (i.e., a minimum of 2 modified reads supporting that site). A nucleotide in a read is defined as ‘modified’ if the modification probability is equal or greater than 0.5 (shown as ‘predicted m^6^A sites’) at the bottom of the IGV snapshot.

An alternative approach to achieve transcriptome-wide RNA modification detection involves *de novo* basecalling of these modifications from the underlying raw signals. Within this approach, the default RNA basecalling model used in the basecalling step would be supplanted by a modification-aware counterpart (illustrated in **Figure 1A**). For instance, an m^6^A modification-aware basecalling model would predict 5 different bases –A, C, G, U and m^6^A– based on the current intensity information. In contrast, the default RNA basecalling confines its predictions to only four bases – A, C, G, U. Notably, the development of modification-aware basecalling models has remained challenging due to the lack of adequate ‘training data’ to train such models.

Here, we train a modification-aware m^6^A basecalling model, which we refer to as *m^6^ABasecaller*, which can predict m^6^A modifications *de novo,* in single reads, and with single nucleotide resolution with high accuracy and low false positive rates. We demonstrate that *m^6^ABasecaller* can yield transcriptome-wide maps of m^6^A modifications across datasets from diverse species and sequencing devices, while the reads are being sequenced, without the need of a knockout/control condition. Moreover, we show that using *m^6^ABasecaller*, it is possible to collect m^6^A modification information at per-isoform level, obtain reproducible and accurate estimates of m^6^A modification stoichiometry transcriptome-wide, analyse m^6^A modification co-occurrence within reads, and characterise the correlation between m^6^A presence and polyA tail lengths, among other features (**Figure 1B**). Our approach can be applied to other RNA and DNA modification types, thus providing a novel framework to train modification-aware base-calling models that can identify an extended set of RNA and/or DNA modifications within individual molecules, and with single nucleotide resolution.

## RESULTS

### m^6^ABasecaller accurately predicts m^6^A modifications in individual reads both *in vitro* and i*n vivo*

Current methods to detect RNA modifications in DRS datasets rely on increased base-calling ‘errors’ and/or altered current intensities at the RNA modified sites. Whilst these methods have proven useful to detect diverse types of RNA modifications ^38,44–54,59,60^, they suffer from important caveats: (i) they require extensive manipulation and processing of the reads before being able to detect RNA modifications (basecalling, mapping, feature extraction, aggregation of features per-site, resquiggling and/or statistical testing) ^61^, ii) they suffer from stoichiometry biases (unmodified reads are preferentially resquiggled) ^38^; iii) they require minimum coverage of ∼30–50 reads to detect a site as modified ^59^; iv) they often require a minimum modification stoichiometry (∼10-20%) to detect a site as modified ^59^, and (iv) the final predictions provided by the different softwares do not provide predictions for individual reads –even if they use per-read information–, but rather, the resolution is at per-site level ^49^.

To overcome these limitations, here we built a novel modification-aware basecalling model that would predict m^6^A modifications *de novo* from raw FAST5 reads, at per-nucleotide and per-read resolution (**Figure 1A**). This model is capable of base-calling 5 different bases (A, C, G, U and m^6^A) in individual reads, completely *de novo*, with single nucleotide resolution, without the need of paired conditions, without the need of minimum coverage per site, and without the need of minimum stoichiometry per-site (**Figure 1B**), thus offering the possibility to study the function and dynamics of m^6^A modifications with an unprecedented resolution.

We first examined the performance of the *m^6^ABasecaller* in synthetic ‘curlcake’ constructs that were either unmodified (0% m^6^A) or fully modified (100% m^6^A) ^65^. Our results showed that *m^6^ABasecaller* predicted m^6^A sites in biologically relevant DRACH sequence contexts with high accuracy and reproducibility (**Figure 3A**, see also **Figure S1**), and with very low amounts of false positives (only 0.3-0.5% were predicted as modified in the same exact sequence context and sites in the unmodified curlcake control sequences).

We then examined the performance of the *m^6^ABasecaller* in *in vivo* datasets. To this end, we ran the *m^6^ABasecaller* on publicly available HEK293T wild type (WT) DRS datasets, as well as on HEK293T METTL3 knockout (KO) and in vitro transcribed (IVT) whole transcriptome human DRS datasets as negative controls (**Figure 3A**, right panel, see also **Table S1** for full list of DRS datasets used in this work). The *m^6^ABasecaller* predicted 6,664 (rep1, 1.2M reads) and 1,625 (rep2, 400K reads) m^6^A-modified sites in the HEK293T WT samples, with a median per-site modification stoichiometry of 15.6 and 15.1%, respectively (**Figure 2B**, see also **Table S2** for number of m^6^A sites predicted by *m^6^ABasecaller*). We observed a sharp decrease in the m^6^A modification levels upon METTL3 KO on the same set of sites, which showed a median per-site modification frequency of 2.8% (rep1) and 2.9% (rep2), respectively (**Figure 2B)**. This decrease in stoichiometry was even more evident in IVT samples, with median modification frequency per-site of 0% (rep1) and 0% (rep2) suggesting that METTL3 KO samples are not completely devoid of m^6^A modifications in their mRNAs, in agreement with recent works ^66^.

**Figure 2.**
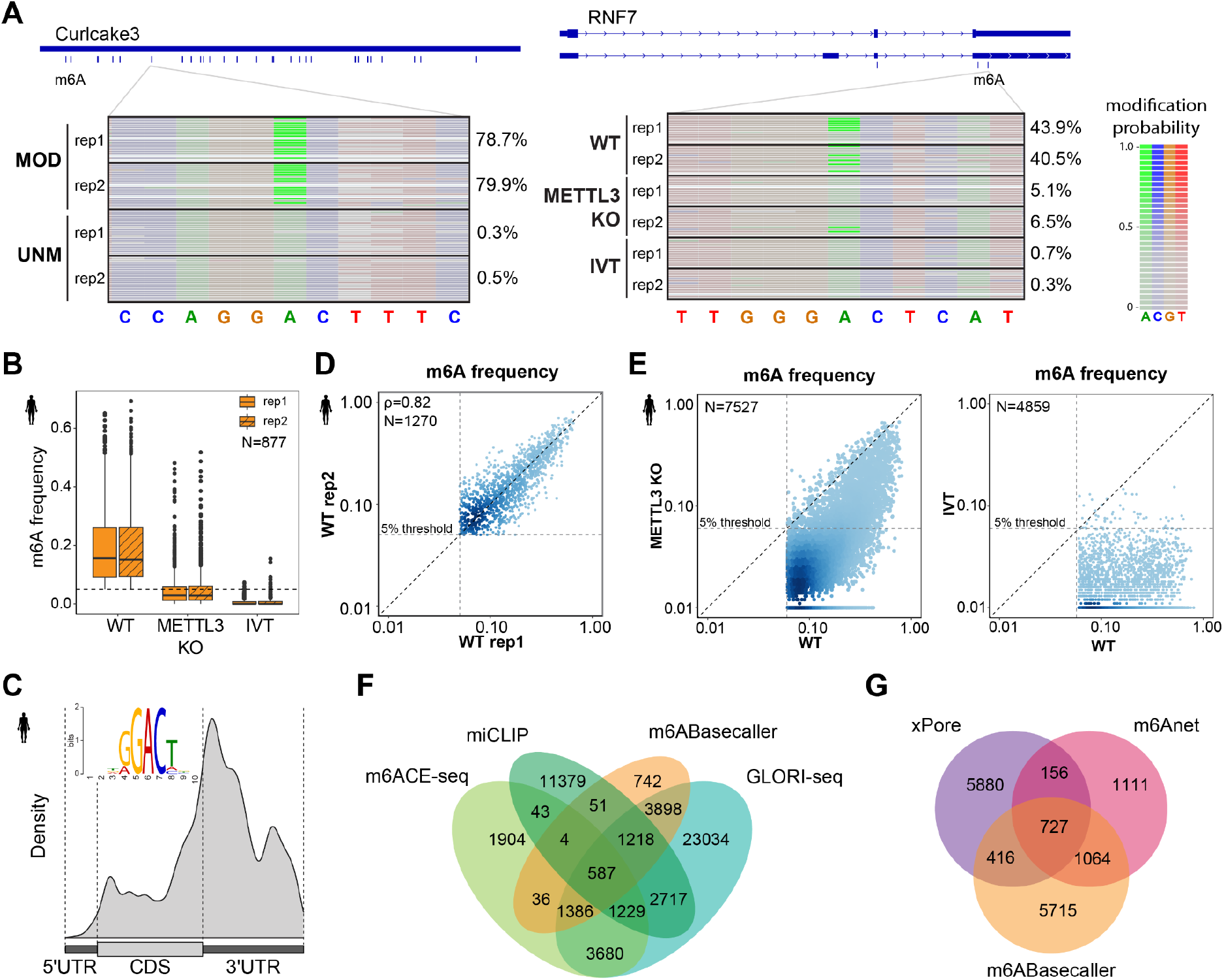
*m^6^ABasecaller* predicts m^6^A in synthetic and native RNA molecules, and shows strong overlap with predicted m^6^A sites using orthogonal methods. **(A)** In the left panel, IGV snapshot of individual reads basec-alled with *m^6^ABasecaller*. The reads are centered at a known m^6^A site, both for synthetic m^6^A-modified curlcake reads (‘MOD’, upper panel) and their unmodified counterpart (UNM, lower panel). In the right panel, reads mapping to human RNF7 gene are shown, in HEK293T WT and METTL3 KO samples, as well as for reads from in vitro transcribed (IVT) human samples. Each row represents a distinct RNA read, and each base from each read has been coloured according to its modification probability. See also **Figure S1** for additional IGV snapshots. **(B)** Boxplot of per-site m^6^A frequencies in two independent replicates of: (i) HEK293T WT, (ii) HEK293T METTL3 KO and (iii) IVT human transcriptome. Only m^6^A sites detected in WT (≥5% m^6^A modification frequency) and with at least 25 reads of coverage in all replicates have been included in this analysis (N=877). The horizontal dashed line indicates the 5% threshold for a site to be identified as “m^6^A-modified”. **(C)** Metagene plot depicting the distribution of m^6^A sites detected in HEK293T WT samples (N=1270). In the upper left corner, the motif obtained with MEME analysing 20 nt sequence context of all replicable sites in HEK293T WT (N=1270) is shown. See also **Figure S2A,B** for metagene plots in additional species. **(D)** Replicability of m^6^A modification frequency in sites with modification frequency greater or equal than 5% and minimum of 25 reads of coverage in both HEK293T WT replicates (Spearman’s R=0.82). Dashed vertical and horizontal lines depict the 5% threshold applied to a m^6^A site to be called. Both axes are log_10_ scaled. **(E)** Scatterplot comparing per-site m^6^A frequencies in modified sites identified in HEK293T cells in WT and upon METTL3 KO (left panel), and in WT compared to IVT control (right panel). Dashed vertical and horizontal lines depict the 5% threshold applied to a m^6^A site to be called. Both axes are log_10_ scaled. (**F**) Overlap between m^6^A sites detected by *m^6^ABasecaller* in HEK293T cells and m^6^A sites predicted using Illumina-based orthogonal methods (m6ACE-seq, miCLIP and GLORI-seq) in HEK293T cells. To provide a comparison across all methods that is independent of sequencing coverage, the set of predicted m^6^A sites by each orthogonal method was reduced to those m^6^A sites for which there was a sufficient coverage in the nanopore DRS dataset, i.e., only m^6^A sites with minimum of 25 reads coverage in the DRS dataset were included in the comparative analysis. (**G**) Comparison of m^6^A sites predicted by *m^6^ABasecaller* and those predicted by other nanopore-based methods (xPore and m6Anet), ran on the same set of reads from HEK293T cells (pooled 2 replicates, see **Table S2**).

**Figure 3.**
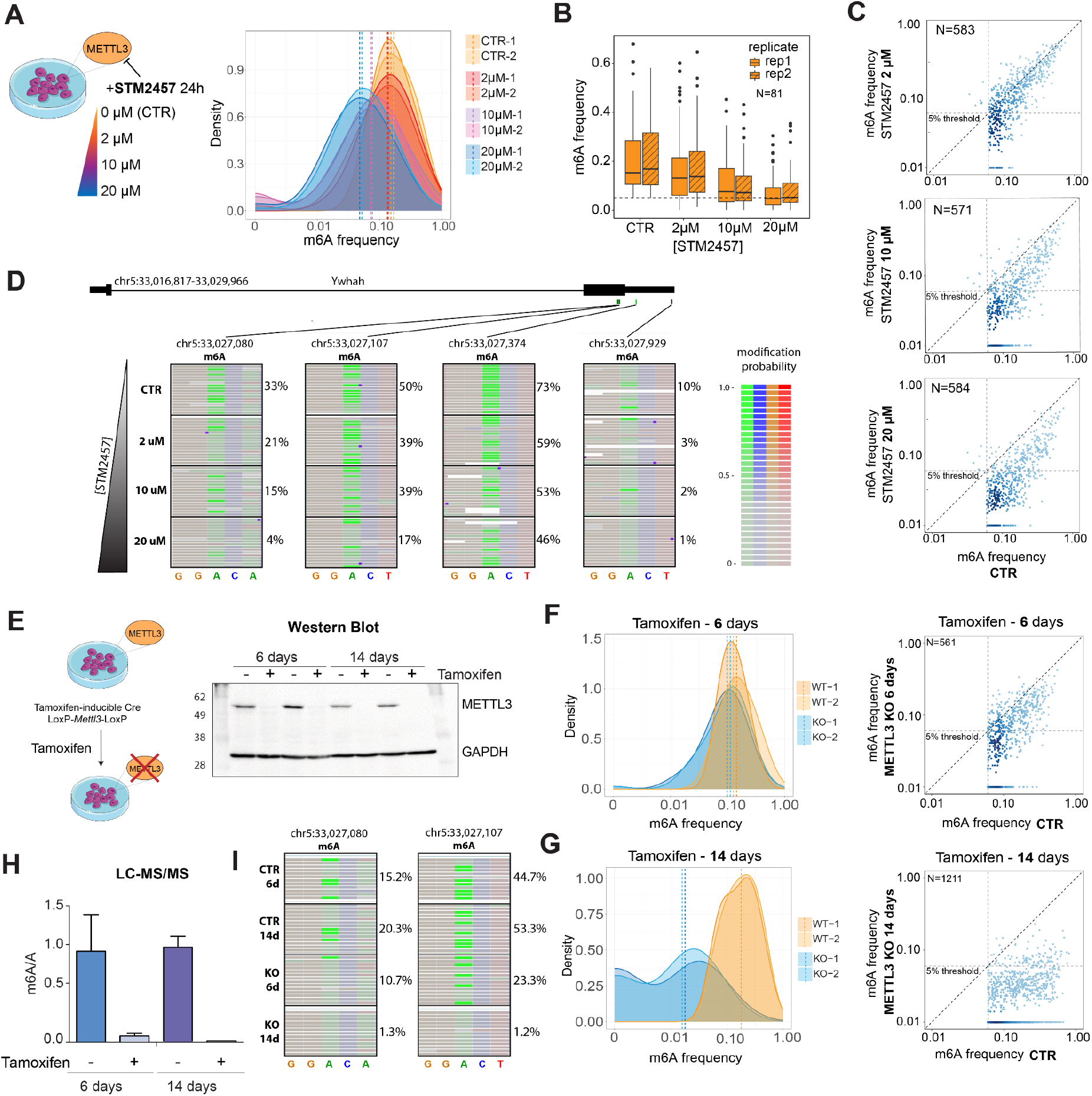
*m^6^ABasecaller* accurately predicts m^6^A modification frequencies transcriptome-wide. **(A)** Density plots of m^6^A modification frequencies in mESC samples treated with different concentrations of STM2457 inhibitor (0, 2, 10 and 20uM). Results are shown for two independent biological replicates. Dashed vertical lines indicate the median m^6^A frequency of each sample. **(B)** Boxplot of the Distribution of the m^6^A frequency at different concentrations of STM2457 in sites with more than 25 reads of coverage in all replicates of all conditions (N=81). **(C)** Scatterplots depicting the per-site m^6^A modification frequency in untreated samples (CTR) relative to STM2457-treated samples (2 µM, upper panel; 10 µM, middle panel; 20 µM upper panel). A gradient from light to dark blue shows the increase in density of data points in the plot. Dashed diagonal black line indicates the x=y line in frequencies. Grey vertical and horizontal dashed lines indicate the 5% m6A frequency threshold used to identify a site as ‘m^6^A-modified’. Axes are log-scaled. **(D)** IGV snapshot depicting the decrease of m^6^A modified reads with increasing concentration of STM2457. The number of reads containing bright green (high probability of m^6^A) diminishes with the increase of STM2457 dosage. **(E)** On the left side, a scheme depicting the generation of tamoxifen-inducible METTL3 KO mESC cell lines is shown. On the right, a Western Blot image depicting the loss of METTL3 upon tamoxifen treatment for 6 days (2 replicates) and 14 days (2 replicates), compared to MetOH-treated cells (CTR) for 6 and 14 days, respectively. GAPDH was used as loading control. **(F,G)** In the left panels, density plot distribution of the m^6^A frequency in the pooled replicates of mES cells treated with tamoxifen (KO) for 6 days (F) or 14 days (G), compared to those treated with MetOH (CTR). In the right panels, scatterplots depicting the modification frequency at m^6^A sites detected in the pooled control samples (CTR) compared to the corresponding frequency in pooled samples treated with tamoxifen for 6 days (F) or for 14 days (G). A gradient from light to dark blue shows the increase in density of data points in the plot. Dashed diagonal black line indicates the x=y line in frequencies. Grey vertical and horizontal dashed lines indicate the 5% m6A frequency threshold used to identify a site as ‘m^6^A-modified’. Axes are log-scaled. For F and G, a pseudocount of 0.001 was added to all values to allow logarithmic scaling of the values. **(H)** Quantification of m^6^A levels in polyA+ RNA from mESC cells treated with tamoxifen for 6 days or 14 days. m^6^A/A is computed as the ratio of m^6^A area vs A area in LC-MS/MS results. 6-day and 14-day tamoxifen treatment led to a ∼15X and 90X reduction in m^6^A levels compared to untreated control samples, respectively **(I)** IGV snapshot depicting the decrease of m^6^A modified reads with increasing duration of tamoxifen treatment. The number of reads containing bright green (high probability of m^6^A) diminishes with the METTL3 inhibition and longer tamoxifen exposure.

We then examined whether the m^6^A sites predicted by the *m^6^ABasecaller* were replicable in independent biological replicates that were independently sequenced, in both mouse and human samples, both qualitatively and quantitatively. Our analyses showed that the overlap of m^6^A-modified sites predicted by the *m^6^ABasecaller* across biological replicates was very high (83-89% overlap, see **Figure S2C,D**). Similarly, the per-site predicted modification stoichiometry was highly replicable across biological replicates (Spearman’s rho=0.82-0.87, see **Figure 2D** and **S2E**). We should note that the correlation of m^6^A modification frequencies across biological replicates improved with increased sequencing coverage, as well as when only m^6^A sites with higher read coverage were included in the analysis (**Figure S2F**).

To further validate the *m^6^ABasecaller*, we examined whether knockout (KO) of METTL3 or METTL14 led to a loss of m^6^A sites identified in the WT samples. Comparative analysis of WT and METTL3 KO samples revealed that 7,369 (98%) and 4,114 (89%) m^6^A sites showed decreased m^6^A frequencies upon METTL3 KO in HEK293T and mES cells, respectively (**Figure 2E**, see also **Figure S3A,B**). Moreover, the median per-site m^6^A frequency decreased from 15.1-15.6% to 2.8-2.9% upon METTL3 KO in HEK293T cells (**Figure S3C,** see also **Table S2**) and from 16.4-18.8% to 4.7-7.8% upon METTL3 and METTL14 KO, respectively, in mES cells (**Figure S3D,** see also **Table S2**).

### m^6^A sites predicted by *m^6^ABasecaller* are largely supported by orthogonal methods

We then assessed whether the m^6^A sites predicted by *m^6^ABasecaller* were also identified using orthogonal methods such as GLORI-seq ^29^, miCLIP ^30^ and m6ACE-seq ^45^. To this end, we examined the overlap between m^6^A sites predicted by *m^6^ABasecaller* in HEK293T cells and those predicted by other orthogonal techniques in the same cell line. Only m^6^A sites for which we had sufficient coverage (>25 reads coverage) in the HEK293T DRS dataset were kept for downstream comparisons: 37,749 sites for GLORI-seq, 17,228 sites for miCLIP, and 8,869 sites for m6ACE-seq (**Table S6**, see also *Methods*). These sites were then compared to the 7,922 sites detected by *m^6^ABasecaller*. Our analyses revealed that 91% of m^6^A sites predicted by the *m^6^ABasecaller* were also predicted by one or more orthogonal methods (**Figure 2F**). More specifically, 89% of the sites identified by *m^6^ABasecaller* were also identified by GLORI-seq (**Figure S4A**), and a more modest overlap (25% and 23%) was observed with m6ACE-seq and miCLIP, respectively (**Figure S4B,C**). We should note, however, that the overlap of sites between Illumina methods themselves was also low –2.8% of predicted m^6^A sites were detected by all 3 Illumina-based methods– (**Figure S4D**).

Finally, we examined the overlap between m^6^A sites predicted by *m^6^ABasecaller* and those predicted by other Nanopore tools, namely xPore ^45^ and m6Anet ^49^, on the same publicly available HEK293T DRS dataset ^45^. Our results showed that only 14% and 22% of the sites detected by *m^6^ABasecaller* were also identified using xPore and m6Anet, respectively (**Figure 2G**). We should note, however, that m6Anet predicted a much lower number of predicted m^6^A sites (3,058) compared to *m^6^ABasecaller* (7,922), possibly due to a minimum modification stoichiometry required to identify a site as ‘modified’ –detection of m^6^A modifications in xPore and m6Anet is per-site, not per-read–. Notably, 58% of the sites predicted by m6Anet were also predicted by *m^6^ABasecaller*.

### *m^6^ABasecaller* recapitulates modification stoichiometry variations upon STM2457 treatment

We then assessed the ability of the *m^6^ABasecaller* to capture variations in m^6^A modification frequencies in biological samples in a quantitative manner. To this end, we sequenced polyA-selected RNA from mESC treated with increasing concentrations of STM2457 (0, 2, 10 and 20 µM), a well-characterised inhibitor of METTL3 ^68^. Our analysis revealed that the m**^6^**A modification frequency observed transcriptome-wide progressively decreased with increasing concentrations of the inhibitor (**Figure 3A**). Indeed, untreated samples showed 15.2-16.7% median m^6^A modification frequency, and decreased to 13.0-13.6%, 7.5-7.1% and 4.7-5.1% upon 2, 10 and 20 µM of STM2457 treatment, respectively (**Figure 3B-D**, see also **Table S2**). Notably, the per-site m^6^A modification stoichiometry values were highly replicable across biological replicates (**Figure S5A,B**), with predicted m^6^A sites matching the DRACH motif (**Figure S5C**), mostly located in the stop codon and 3’UTR regions of coding transcripts (**Figure S5D**), in agreement with previous reports ^26,27,67^. Overall, our results show that the *m^6^ABasecaller* can robustly identify quantitative changes in m^6^A modification stoichiometry in a transcriptome-wide fashion and in a replicable manner, in addition to qualitative changes transcriptome-wide.

### m^6^ABasecaller reveals incomplete loss of m^6^A in inducible METTL3 KO systems

To further validate the *m^6^ABasecaller*, we generated an inducible METTL3 KO cell line by introducing 2 LoxP sequences upstream of the METTL3 exon 2 and downstream of the METTL3 exon 4 –as described in ^69^– into a mER-Cre-mER mESC line, which constitutively expresses Cre recombinase fused to a mER domain, inducing its translocation to the nucleus upon tamoxifen treatment, thus making the knockout system tamoxifen-inducible (**Figure 3E**, see also *Methods*). To determine the duration of tamoxifen treatment required to observe a complete loss of METTL3 protein, mESC were treated with tamoxifen for 1, 3, 6 and 14 days, and METTL3 protein levels were quantified using Wesrwen Blotting, showing that 6 and 14 days –but not 1 or 3 days– of tamoxifen treatment led to a complete loss of METTL3 protein in all examined clones (**Figure 3E**). Thus, all subsequent analyses were performed using 6-day tamoxifen-treated (which we refer to as ‘METTL3 KO’) or vehicle-treated (‘CTR’) mES cells.

We then sequenced METTL3 KO and CTR mESC cells in biological duplicates using nanopore DRS (**Figure S6A**), and predicted m^6^A-modified sites in individual reads using *m^6^ABasecaller*. To our surprise, we only observed a modest reduction in the median m**^6^**A modification frequencies when comparing CTR (∼13% median m^6^A frequency) and METTL3 KO samples (∼10.5% median m^6^A frequency) (**Figure 3F**, see also **Table S2**), Moreover, only 41% of m^6^A sites identified in CTR samples (n=231) disappeared upon tamoxifen treatment –i.e., fell below the 5% modification frequency threshold– despite these conditions leading to a complete loss of METTL3 protein (**Figure 3E**).

Puzzled by these results, we hypothesised that the loss of METTL3 might not lead to an immediate loss of m^6^A modifications in mRNA molecules, and that our observations could be explained by the presence of m^6^A modifications that were previously deposited in mRNAs that have not yet been degraded. We reasoned that if this were the case, the 14-day tamoxifen-treated samples should show significantly less m^6^A than the 6-day tamoxifen-treated cells. To test this, we sequenced polyA-selected RNA from 14-day tamoxifen-treated and matched CTR mESC cells in biological duplicates (**Figure S6B**) and analysed the DRS data using *m^6^ABasecaller*, using same settings and parameters than previously employed for 6-day treated/untreated samples. Notably, this time we observed a drastic reduction in the median m**^6^**A modification frequencies when comparing untreated (∼18% median m^6^A frequency) and 14-day tamoxifen-treated samples (∼1.7% median m^6^A frequency) (**Figure 3G**, see also **Table S2**). Indeed, 91% of m^6^A sites identified in CTR samples (n=1,106) falling below the 5% modification frequency detection threshold upon tamoxifen treatment. Thus, our results support the hypothesis that there is a delay between the loss of METTL3 protein and the loss of m^6^A in mRNAs.

To further validate the results obtained using *m^6^ABasecaller*, we examined the m^6^A modification levels in polyA-selected RNAs from 6- and 14-days tamoxifen-treated samples using Liquid Chromatography coupled to Mass Spectrometry (LC-MS/MS)^70^. This analysis revealed a 14-fold decrease in m^6^A levels in 6-day tamoxifen-treated samples, compared to CTR untreated samples (**Figure 3H**). This difference was further increased upon extending the tamoxifen treatment; 14-day tamoxifen-treated showed a 90-fold decrease in m^6^A levels, compared to 14-day CTR untreated samples. Overall, the LC-MS/MS results support our model, and are in agreement with the results observed by *m^6^ABasecaller*, demonstrating that while m^6^A levels globally decrease upon induced METTL3 protein loss, absence of METTL3 protein does not guarantee absence of m6A in mRNAs, as some m^6^A-modified mRNAs might require additional time to disappear (**Figure 3I**).

### Per-read analysis reveals direct correlation between m^6^A, polyA tailing and splicing

A major strength of the *m^6^ABasecaller*, compared to Illumina-based methods and other nanopore-based methods that do not have per-read resolution, is that in addition to providing m^6^A modification information at single nucleotide and single molecule resolution, it allows direct correlation between the presence of m^6^A and other post-transcriptional features that can be identified and/or measured in the same RNA molecules, such as polyadenylation or splicing. Indeed, *m^6^ABasecaller* allows investigating questions such as: i) whether m^6^A modifications co-occur simultaneously in the same read or are mutually exclusive (‘m^6^A co-occurrence’); ii) whether the presence of m^6^A affects polyA tail lengths; and iii) whether m^6^A modifications are (un)equally deposited across isoforms of a same gene (**Figure 4A**).

**Figure 4.**
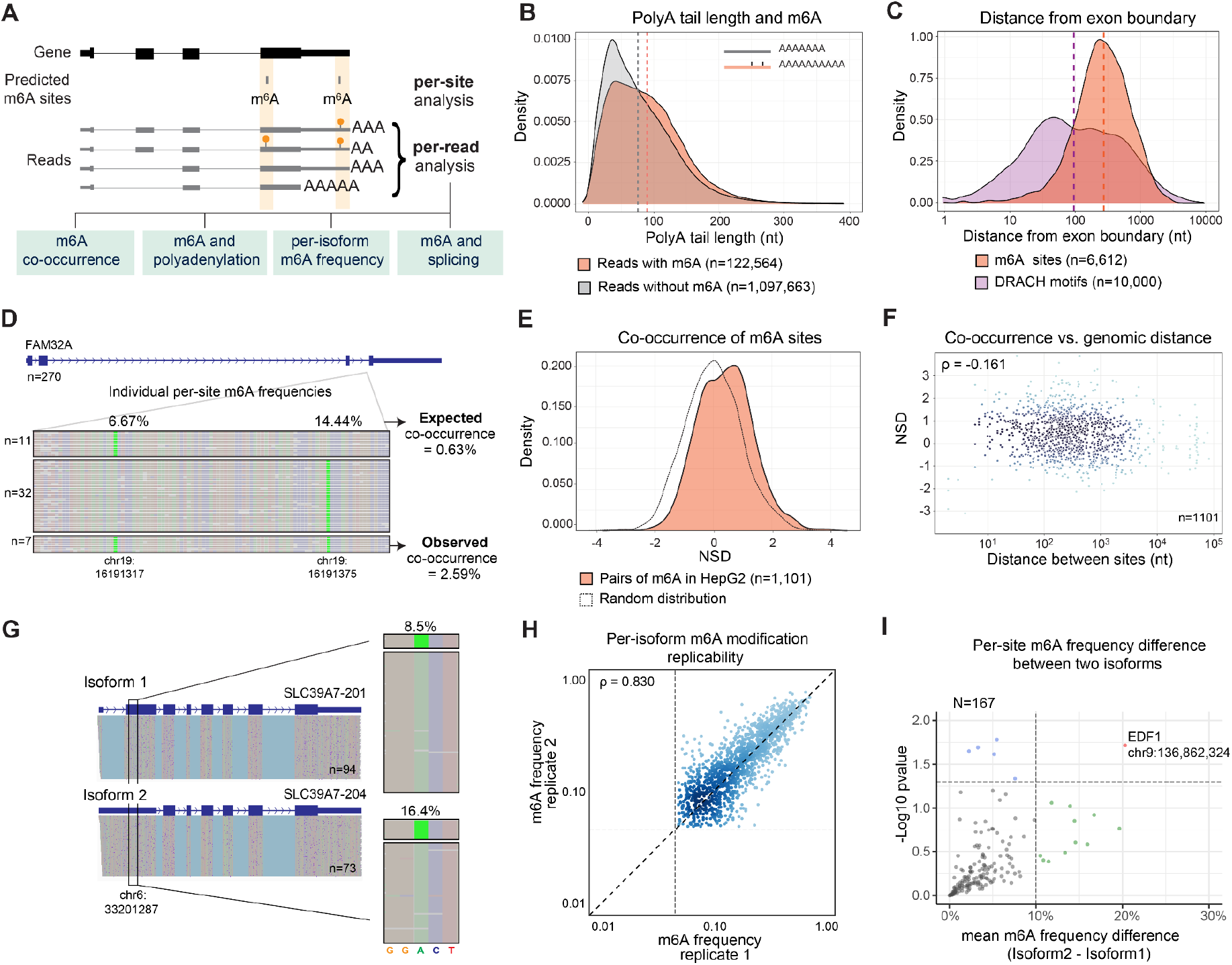
Analysis of m^6^A modifications at per-read level. **(A)** Schematic overview of the distinct layers of information that can be studied with per-read resolution. **(B)** Distribution of polyA tail lengths in reads containing at least 1 m^6^A site (orange) and reads without m^6^A (grey). Dashed line indicates the median polyA tail length of each group. **(C)** Distribution of the distance between each m^6^A modification and the closest boundary of an exon (orange), compared to the distribution of the same distance calculated for a random subset of DRACH motifs in the same genes (purple). The x axis is log_10_-scaled. **(D)** IGV snapshot of reads mapping to FAM32A gene. Bases have been coloured according to modification probability, showing that this gene contains two m^6^A modifications at positions chr19:16191317 and chr19:16191375, depicted with bright green colour. Reads have been binned depending on whether they contain one m^6^A modified site (chr19:16191317), the other m^6^A modified site (chr19:16191375), or both sites modified. The observed and expected co-occurrence values, given the individual per-site m6A modification frequencies, are also shown. Reads that had both positions unmodified are not shown. **(E)** Distribution of NSD values, which quantifies the co-occurrence of pairs of m^6^A sites (n=1,101) in HepG2 cells is shown in red. As a control, a random distribution generated with the same amount of data points and the same standard deviation, centered in 0, is also shown. **(F)** For each pair of m^6^A sites (n=1,101) analysed, the NSD is plotted against the genomic distance (log_10_-scaled) between the two sites. Each dot represents a pair of m^6^A sites. Spearman correlation is shown. **(G)** IGV snapshot depicting the modification frequency at position chr6:33201287 in both isoforms (SLC39A7-201 and SLC39A7-204) from gene SLC39A7. Modification frequency at per-isoform level is shown. **(H)** Scatterplot depicting the correlation of m6A frequencies at per-isoform level. Grey vertical and horizontal dashed lines indicate the 5% m^6^A frequency threshold used to identify a site as ‘m^6^A-modified’. Axes are log-scaled. **(I)** One-sided volcano plot depicting isoform-specific m^6^A modification patterns. In the y axis, the mean absolute difference in ‘m^6^A modification frequencies across 2 isoforms (n=167 comparisons) in two replicates is shown. To increase the statistical power of the analysis as well as the number genes for which isoform-specific m^6^A analysis was possible, all HepG2 MinION reads were pooled as one replicate (n=4,741,372 reads, see **Table S1**). HepG2 reads from a PromethION run were used as a second replicate (n=6,200,572). Only reads unambiguously assigned to a given isoform were kept for the analysis.

To address these questions, we processed publicly available DRS runs from HepG2 polyA-selected RNA, generated by the Singapore consortium ^71^. These runs included more than 13 million reads (**Table S1**), thus making it possible to perform per-isoform m^6^A analyses on this dataset. The *m^6^ABasecaller* identified a total of 28,846 m^6^A sites (listed in **Table S7**) in the pooled set of 13 million HepG2 reads. Mapped reads were then filtered to retain only full-length reads that were uniquely assigned to one isoform, which were subsequently used for per-isoform m^6^A analyses detailed below.

Firstly, we examined whether the presence of m^6^A in mRNAs globally affected polyA tail lengths. To this end, RNA molecules were binned into two groups: (i) those with one or more m^6^A modifications, and (ii) those without m^6^A modifications. We compared the polyA tail length distributions of the two groups, finding that m^6^A-containing mRNA molecules showed significantly longer polyA tail lengths than their unmodified mRNA counterparts (Mann-Whitney-Wilcoxon test p<2.2e^-16^, **Figure 4B**). Moreover, we found that the increase in polyA tail length was directly correlated with the number of m^6^A sites present in the RNA molecule (**Figure S7A**).

Recent works have provided evidence for m^6^A being excluded from exon junctions ^72–74^. Thus, we wondered whether the same trend would be observed in DRS datasets analysed with *m^6^ABasecaller*, as they provide both isoform and m^6^A information for each RNA molecule sequenced. To this end, we computed the distance between each m^6^A site (n=6,612 sites, those that could be unequivocally assigned to a single transcript), to the closest exon start or end. As a control, the distance between randomly-selected DRACH motifs and the closest exon start or end was also computed (see *Methods*). Our analysis revealed that the distance between m^6^A sites and the closest exon boundary was significantly higher (median=274 nt) than the one expected by chance in randomly chosen DRACH motifs (median=95 nt) (p-value=1.97e^-12^, see also **Figure 4C**), in agreement with previous observations ^72–74^ using orthogonal methods.

### m^6^A modifications are preferentially deposited on the same mRNA molecule

mRNA molecules are typically substochiometrically modified, implying that not all molecules mapping to a given m^6^A site are m^6^A-modified. Whether m^6^A modifications are preferentially deposited in a subset of mRNA molecules (‘hyper-modified RNAs’) or uniformly placed across molecules is largely unknown, mainly due to our inability to map m^6^A modifications in individual RNA molecules. Here we exploited the single molecule resolution feature of the *m^6^ABasecaller* to address this question. To this end, we first extracted m^6^A information from full-length RNA reads that could be unambiguously assigned to a given mRNA isoform (see *Methods*). For each pair of m^6^A-modified sites present in a given isoform, we computed the ‘expected co-occurrence’ by multiplying the m^6^A modification frequencies of each of the two m^6^A-modified sites (n=1,101 pairs of m^6^A sites in the HepG2 dataset). Then, we compared this value to the ‘observed co-occurrence’ (**Figure 4D**, see also **Figure S8A**), which we defined as the proportion of molecules that have both m^6^A sites modified. Finally, for each pair of m^6^A sites, we calculated the number of standard deviations (NSD) that the ‘observed co-occurrence’ deviated from the expected co-occurrence (see *Methods*). Therefore, pairs of sites with NSD>0 co-occur more frequently than expected, whereas m^6^A pairs with NSD<0 are preferentially deposited in distinct RNA molecules.

We examined the distribution of NSD values for all pairs of m^6^A sites transcriptome-wide, finding that the distribution of NSD values was slightly shifted towards positive values (**Figure 4E**). To assess whether this shift was significant, we compared this distribution to a random distribution with same sample size and variance, but mean value of 0, finding that the two distributions were significantly different (Mann-Whitney, p=3.20e^-20^). We also examined the correlation between co-occurence of m^6^A sites and the genomic distance between the set of m^6^A pairs analyzed, finding no positive correlation (Spearman’s ρ=-0.161) (**Figure 4F**). Thus, our results suggest that m^6^A modifications preferentially co-occur in the same RNA molecules, independently of the distance between the two sites, implying that the deposition of an m^6^A modification in a given mRNA molecule is more likely to occur on mRNA molecules with a previous m^6^A modification.

### m^6^A stoichiometry does not largely vary across isoforms

We then wondered whether m^6^A modification stoichiometries of a given m^6^A site were similar or distinct across isoforms, as we could eyeball some genes that apparently showed different m^6^A modification stoichiometries across isoforms (**Figure 4G**). To address this question systematically and in a transcriptome-wide fashion, we used high-coverage HepG2 human datasets (**Table S2**). Firstly, we examined whether per-isoform m^6^A stoichiometries were replicable across independent biological replicates, finding an overall Spearman’s correlation of 0.83 between biological replicates sequenced in different flowcells (**Figure 4H**, see also *Methods*). We then examined whether the modification stoichiometry of m^6^A sites varied consistently across isoforms. To this end, we analysed the variation in isoform-specific modification stoichiometry of m^6^A sites that showed a minimum per-isoform coverage of 40 reads in both replicates (n=167 sites). Analysis of differential modification levels revealed only modest differences in m^6^A modification stoichiometries across isoforms (**Figure 4I**), suggesting that m^6^A stoichiometry is in general not isoform-specific. Indeed, most of the identified m^6^A changes across isoforms were not replicable across biological replicates, suggesting that the initial variations observed across isoforms were caused by random sampling differences. The only site that we found to be significant was found in the EDF1 gene, which showed a replicable 20% stoichiometry difference between two isoforms (**Figure S8B**). Notably, the lowest m^6^A frequency was found in the isoform that had a splicing site ∼20nt downstream from the m6A-modified site.

## DISCUSSION

Nanopore sequencing technologies are revolutionising the fields of genomics and transcriptomics. Despite their potential to improve the precision, quality and complexity of existing DNA and RNA modification maps, long-read sequencing methodologies have still not been adopted as a mainstream sequencing technology. A major obstacle for their widespread adoption stems from the lack of modification-aware base-calling algorithms that will work for any given sequence context. To date, efforts to detect modifications *de novo* via base-calling models have primarily been focused on DNA modifications: species- and/or context-specific modification-aware basecalling models are available for 4mC, 5mC, 5hmC and 6mA. By contrast, modification-aware RNA basecalling models are currently unavailable, with the exception of a few models released by the community that were trained on a small subset of synthetic sequences, rendering them inapplicable for transcriptome-wide analyses ^62,75^.

In the last few years, several works have successfully shown that m^6^A RNA modifications –as well as other RNA modifications– can be detected using nanopore sequencing ^44–51^. However, methods developed so far lack single molecule resolution (can only provide m^6^A predictions at per-site level, not per-read), require computationally-intensive steps in which they analyse aggregated per-read information and have relatively high false positive and false negative rates ^50^. Here we address these limitations with a novel approach that provides reliable m^6^A information at per-read level, the *m^6^ABasecaller* (**Figure 1**), which allows us to address questions regarding the mechanism of m^6^A deposition in mRNAs or the interplay between m^6^A modifications and polyA tail lengths (**Figure 4**). In contrast to previous methods, the *m^6^ABasecaller* does not require a control sample (knockout or unmodified) to perform its predictions, nor a minimum sequencing coverage to predict a nucleotide as m^6^A-modified; the prediction is performed per-read and per-nucleotide, independently of other reads, m^6^A sites, or datasets.

The *m^6^ABasecaller* opens new doors to study the dynamics of m^6^A modifications across tissues, cell types and conditions at an unprecedented resolution, as well as the interplay between different regulatory layers at per-read resolution. Notably, the *m^6^ABasecaller* has also allowed us to identify unexpected issues that may arise when using inducible knockout (iKO) or knock-down (KD) systems as ‘control’ conditions. Indeed, we find that most m^6^A-modified sites are still present at detectable levels upon shorter induction times (**Figure 3E-F**, see also **Figure S3**) suggesting that caution should be taken when using these systems and interpreting their results. Moreover, our work demonstrates that the loss of METTL3 protein in inducible systems is not sufficient to prove that m^6^A is absent in mRNAs, and that, even if at lower stoichiometries, m^6^A modifications will still be present in certain RNA molecules.

Overall, *m^6^ABasecaller* is a fast and robust tool that allows us for the first time to produce transcriptome-wide m^6^A maps at single molecule and single nucleotide resolution, in full-length mRNA reads, with high specificity and high sensitivity (**Figure 2**), opening its use for a variety of applications, including the direct detection of m^6^A modifications in clinical samples using nanopore sequencing.

## MATERIALS AND METHODS

### mESC culturing and passaging

*M. musculus* embryonic stem cells (mESC E14tg2A) were cultured under feeder-free conditions using 0.1% Gelatine (Millipore, #ES-006-B) coated plates (ThermoFisher, #140675) and grown in mESC medium prepared as follows: KnockOut™ DMEM (ThermoFisher, 10829018) supplemented with 10% FBS (in-house tested for ES competence) MEM Non-Essential Amino Acids Solution 1X (ThermoFisher, #11140050), GlutaMAX™ 2 mM (ThermoFisher, #35050061), Pen/Strep 1X (ThermoFisher, #15140122), β-mercaptoethanol 50 µM (Gibco, #31350010), HEPES 30 mM (Gibco, #15630080), 0,22 µm filtered and then supplemented with LIF conditioned (may 2021- in house tested). Cells were passed every 2-3 days in a 1:6-1:8 dilution.

### Generation of tamoxifen-inducible METTL3 knockout mES cells

In order to obtain tamoxifen-inducible METTL3 Knock-Out mES cells, 10^6 E14 mES (mESC E14tg2A) cells were transformed via electroporation (EP) with 3 ug of Piggybac MerCreMer Addgene plasmid (#124183) and 9ug pBase plasmid (PL623 - Wellcome Trust Sanger Institute) in NEPA21 electroporator. Electroporated (EP) cells were then selected with 400 ug/mL Neomycin (Roche, #04727878001) and resistant clones (mES cell line expressing Cre) were established. Subsequently, cells carrying the MerCreMer system were used for the insertion of 2 LoxP sites flanking a fragment of METTL3 gene. Briefly, the LoxP1 site was inserted by electroporation of 12,2 uM RNP Cas9 protein (PNA BIO, #CP02), 16 uM sgRNA1 and 4,8 uM ssODN (LoxP1 sequence) in OptiMEM (Gibco, #11058-021). 24h post-EP the cells were sorted, seeded at single cell and genotyped by PCR and Sanger sequencing. One positive clone was then selected for the insertion of the LoxP2 sequence using the same procedure as described above with sgRNA2. 48h post-EP, the cells were sorted, seeded at single cell and genotyped by PCR and Sanger sequencing. All sequences used to generate this cell line can be found in **Table S8**, and were taken from Wang et al., 2018 ^69^.

### Treatment of mESC with STM2457 inhibitor and with Tamoxifen

The METTL3 inhibitor STM2457 (STORM Therapeutics) was administered to mESC cultures to a final concentration of 2, 10 or 20 µM. Dilutions of STM2457 were performed in DMSO (PanReac AppliChem, #A3672). As a control, the same volume of DMSO was administered to the cells. Cells were harvested with Trizol (Invitrogen,15596018) 24h after the addition of the inhibitor. For tamoxifen treatment, (Z)-4-Hydroxytamoxifen (Sigma, #H7904-5MG) (diluted in MetOH) was added to the culturing media to a final concentration of 2.5 µM at each medium replacement until harvest day. As a control, we added the same volume of vehicle (MetOH). Cells were harvested in Trizol (Invitrogen, #15596018) 6 or 14 days after tamoxifen addition to the media.

### Western Blotting

To perform the Western Blot assay, cells were lysed in RIPA buffer containing protease inhibitor (Roche, #11873580001) and lysates were centrifuged at 15000 rcf for 15 min at 4°C to remove cellular debris. Lysates were then prepared with NuPAGE sample buffer LDS 4X (Thermo Fisher, #NP0007) and NuPAGE Sample Reducing Agent (Thermo Fisher, #NP0004), and were loaded on 15% acrylamide gels. Protein content was previously checked with Ponceau (Sigma-Aldrich, #P7170) and membranes (Merck, #IPVH00010) were blocked with TBS-T BSA 3%. METTL3 antibody (Abcam ab195352) was used at a 1:1000 dilution, and GAPDH antibody (Abcam ab8245) was used at 1:10.000. Anti-mouse (Agilent #P0260) or anti-rabbit (Abcam, #ab6721) HRP-conjugated secondary antibodies were diluted 1:5000 in TBS-T.

### Total RNA extraction

For each condition, 3 wells of a 6-well plate of mESC (corresponding to 6*10^6 cells approximately) were resuspended in 1 mL Trizol (Invitrogen,15596018) vortexed two times for 30 seconds and incubated 5 minutes at room temperature, then processed immediately or stored at −80. Next, 200 µL of chloroform (Sigma, C2432) was added to each sample, mixed for 20 seconds, incubated for 2-3 minutes at room temperature and centrifuged at 16,000 g for 15 minutes at 4°C. The resulting upper aqueous phase was transferred to a new tube and mixed with 500 µL of molecular grade 2-propanol (Sigma, I9516). Samples were incubated 10 minutes at room temperature and centrifuged at 12,000 g for 10 minutes at 4°C to precipitate the RNA. The pellet was washed with 70% ethanol and centrifuged at 7,500 g for 5 minutes at 4°C, air-dried for 10 minutes and eluted in nuclease free water.

### PolyA selection from mESC total RNA

Total RNA was DNAse-treated (Ambion, AM2239) at 37°C for 20 minutes, and cleaned up using RNeasy MinElute Cleanup Kit (Qiagen, 74204). 70-100 ug of total RNA was then subjected to double polyA-selection using Dynabeads Oligo(dT)25 (Invitrogen, 61002) following manufacturer’s protocol and eluted in ice-cold 10 mM Tris pH 7.5.

### Direct RNA nanopore library preparation and sequencing

PolyA(+) selected RNA samples were prepared for nanopore sequencing using the direct RNA sequencing (DRS) kit (SQK-RNA002), following the ONT protocol version DRS_9080_v2_revI_14Aug2019 with half reaction for each library until the RNA Adapter (RMX) ligation step, with some adaptations. Briefly, 250ng of polyA(+) RNA were ligated to pre-annealed custom RT adaptors (IDT) containing barcodes ^78^ with T4 DNA concentrated Ligase (NEB-M0202M) for 15 min at RT. Next, reverse transcription was performed during 15 min at 50°C using SuperScript IV RT Enzyme (Invitrogen,18090050), followed by heat inactivation for 5 minutes at 70°C. Ligated products were then purified using 1.8X Agencourt RNAClean XP beads (Fisher Scientific, NC0068576) and washed with 70% freshly prepared ethanol. For the last ligation step, 50 ng of reverse transcribed RNA from each reaction was pooled, mixed with RMX adapter, composed of sequencing adapters with motor protein, and incubated for 15 minutes in the presence of concentrated T4 DNA Ligase (NEB-M0202M). Finally the ligated RNA:DNA hybrid was purified using 1X Agencourt RNAClean XP beads, washed with Wash Buffer (WSB) twice. The sample was then eluted in Elution Buffer (EB) and mixed with RNA Running Buffer (RRB) before loading onto a primed R9.4.1 flowcell, and ran on a MinION sequencer.

### Basecalling, demultiplexing direct RNA sequencing data

Raw Fast5 files from dRNA sequencing runs analysed in this study (listed in **Table S1**) were processed with Master of Pores pipeline version 2 ^61^, which is publicly available in GitHub (https://github.com/biocorecrg/MOP2). The *mop_preprocess* module was used to process the samples, using *DeePlexiCon* with default parameters ^78^ to demultiplex the runs when required, and basecalled with *Guppy* basecaller 3.4.5 (https://nanoporetech.com) using the m^6^A basecalling model trained as part of this work (*rna_r9.4.1_70bps_m6A_hac.cfg*, which is available at https://github.com/novoalab/m6ABasecaller/tree/main/basecalling_model).

### Extraction of modification information from DRS datasets basecalled with m^6^ABasecaller

m^6^A-basecalled Fast5 files were processed with ModPhred ^62^ to encode the modification probabilities (calculated by the *m^6^ABasecaller*) into Fastq files. ModPhred was also used to map the Fastq files with minimap2 ^79^ (version 2.17-r941) with “*-ax map-ont -k13”* parameters to the hg38 genome for human data, mm10 genome for mouse data, and GRCz11 for zebrafish data. ModPhred was also used to generate a compressed bedMethyl (*.mod.gz) file with a list of m^6^A sites, which were considered as those positions with coverage>=25 and modification frequency >= 5%.

The output file generated by ModPhred (*mod.gz*) was then processed to identify replicable m^6^A sites, which were defined as those sites with coverage>=25 and their respective modification frequency>= 5% in all replicates. Metagene plots depicting the distribution of m^6^A sites were generated with a custom R script based on the Guitar package version 2.14 ^80^. Motif enrichment analysis of the predicted m^6^A sites was performed using MEME version 4.11.2 ^81^ with the following parameters: *-nostatus -dna -mod zoops -nmotifs 5 -minw 2 -maxw 10*. To build m^6^A frequency scatterplots and density plots with logarithmic axis, a pseudocount of 0.001 was added to each value to allow for logarithmic transformation.The set of m^6^A sites predicted in human, mouse and zebrafish DRS datasets using *m^6^ABasecaller* are listed in **Table S3** (human HEK293T), **Table S4** (mouse ES cells), **Table S5** (zebrafish embryos 4hpf) and **Table S7** (human HepG2).

### Isoform annotation

Per-read isoform annotation was performed on uniquely mapping reads, which were extracted using the samtools flag -F 3844, using Isoquant v2.2.2 ^82^ with “--stranded forward --complete_genedb --count_exons --data_type nanopore” parameters and the Ensembl annotation version 109. Isoquant was used to perform isoform predictions, and only those reads with “unique”, “fsm” (full splice match) or “mono_exon_match” tags were kept for downstream analyses. We should note that this filtering discarded ∼80% of the mapped reads that could not be unambiguously assigned to a specific isoform.

### PolyA tail length estimation

Per-read polyA tail length estimation was performed using the *mop_tail* module of MOP2, using *tailfindr* version 1.3 ^83^. Per-read polyA tail length estimations were merged with per-read modification data and per-read isoform assignments, generating a final table that contained all features for every read.

### Analysis of m^6^A modification co-occurrence

To assess whether m^6^A modifications tend to occur in the same reads, we first filtered the DRS dataset to keep only full-length reads that were uniquely assigned to a given isoform. Then, we analysed the per-read m^6^A modifications for each pair of predicted m^6^A sites that met the following criteria: (i) both m^6^A sites were found in a transcript with coverage >=200 reads, and (ii) the expected number of reads that contain two modified sites >= 2. If these criteria were met, the expected m^6^A co-occurrence of site A and B –quantified as ‘reads’ or ‘counts’– was calculated as follows:

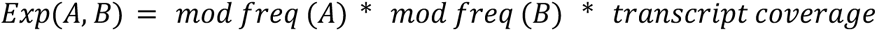

Then, for each pair of m^6^A sites analysed, the number of standard deviations (NSD) from the expected counts was computed as follows:

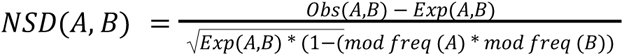

### Comparison of m6ABasecaller predicted m^6^A sites with orthogonal datasets

To compare the list of m^6^A sites predicted by the *m^6^ABasecaller* to those previously published and had been annotated under hg19 assembly, we lifted the genomic coordinates using the UCSC “Lift Genome Annotations” online tool (https://genome.ucsc.edu/cgi-bin/hgLiftOver) to hg38. To subset the bed files based on coverage (only sites with at least 25 reads coverage were included in our analysis), we used bedtools coverage tool (bedtools version v2.30.0). The m^6^A sites identified by *m^6^ABasecaller* as well as by other orthogonal studies (before and after filtering) are listed in **Table S6**.

## DATA AVAILABILITY

Basecalled FAST5 generated as part of this work have been deposited in ENA under the accession number PRJEB61874. The deposited data includes: (i) DRS runs from in vitro transcribed curlcakes (12.5%, 25%, 50% and 75% of m^6^A), (ii) DRS runs from mES cells treated with STM2457 (CTR, 2uM, 10uM and 20 uM); and (iii) DRS runs untreated and tamoxifen-treated (6 days or 14 days) mES cells, where the tamoxifen treatment induces METTL3 KO. All other datasets used in this work were taken from publicly available datasets, and are summarised in **Table S1**. The list of m^6^A-modified sites in HEK293T cells predicted by Illumina-based methods (GLORI-seq and m6ACEseq) used for orthogonal validation were taken from publicly available resources (GLORI-seq from Supplementary Data 1 of ^29^ and m6ACEseq from m6ACE-Seq.csv from ^84^). The list of predicted m^6^A-modified sites in HEK293T cells predicted using miCLIP was kindly provided by Samie Jaffrey. Native and PCR amplified *E. coli* C600 reads were kindly provided by Jan Gawor from the Institute of Biochemistry and Biophysics Polish Academy of Sciences in Warsaw.

## CODE AVAILABILITY

Code to basecall the DRS data using *m^6^ABasecaller* can be found in the *m^6^ABasecaller* GitHub repository (https://github.com/novoalab/m6ABasecaller). The trained *m^6^ABasecaller* model is publicly available in the *m^6^ABasecaller* GitHub repository.

## ACKNOWLEDGEMENTS

We thank all the members of the Novoa lab for their insightful discussions. SC was supported by “la Caixa” InPhINIT PhD fellowship (LCF/BQ/DI19/11730036) and is currently supported by Centro de Excelencia Severo Ochoa funding. AD-T is supported by an FPI Severo-Ochoa fellowship by the Spanish Ministry of Economy, Industry and Competitiveness (MEIC). LPP was supported by funding from the European Union’s H2020 research and innovation programme under Marie Sklodowska-Curie grant agreement No. 754422. This work was supported by funds from the Australian Research Council (DP180103571 to EMN), the Spanish Ministry of Economy, Industry and Competitiveness (MEIC) (PID2021-128193NB-100 to EMN) and the European Research Council (ERC-StG-2021 No 101042103 to EMN). We acknowledge the support of the MEIC to the EMBL partnership, Centro de Excelencia Severo Ochoa and CERCA Programme / Generalitat de Catalunya. We would like to thank the CRG Scientific Information Technologies (SIT) facility for setting up the GPU cluster and continuously upgrading the corresponding software that was needed to benchmark the software using diverse Guppy and CUDA versions. We also thank the CRG Tissue Engineering Facility for their help in establishing the tamoxifen-inducible METTL3 knockout cell lines. We thank STORM Therapeutics for sharing the STM2457 METTL3 inhibitor.

## AUTHOR CONTRIBUTIONS

SC cultured untreated and tamoxifen-treated mESC cells, STM2457-treated mES cells, extracted their RNA, and built their corresponding direct RNA sequencing libraries from the polyA(+)-selected RNA fractions. AD-T developed custom python scripts to analyse the data at per position and read level. SC and AD-T analyzed both in-house produced and publicly available datasets for the evaluation of the performance of the *m^6^ABasecaller*. LPP developed the method for basecaller training (NanoRMS2) and trained all the basecalling models used in this work, including the *m^6^ABasecaller*. RM performed initial benchmarking of the tamoxifen-inducible METTL3 knockout mESC cell lines. LL generated the *in vitro* transcribed m^6^A-modified curlcakes. LPP and EMN conceived the work. EMN supervised the work and acquired funding. SC, AD-T, LPP and EMN wrote the manuscript, with the contribution from all authors. The order of the first two co-first authors was determined randomly by coin flipping.

