## Supplementary Information for "*De novo* basecalling of m^6^A modifications at single molecule and single nucleotide resolution"

### 1. SUPPLEMENTARY FIGURES

**Figure S1. Performance of m6ABasecaller in synthetic m6A modified RNAs. (A)** IGV snapshots of positions predicted as m6A modified in 100% m6A curlcake. Individual reads are reported as “collapsed” and bases are colored by quality score, which contains information about m6A probability. The first two panels show reads from 2 independent m6A-modified curlcakes, and on bottom panels show individual reads from 2 independent unmodified curlcakes. The underlying reference sequence is shown in the bottom of each panel, and the predicted m6A frequency by the m6ABasecaller for each site and replicate is shown in the right of each panel. **(B)** Replicability between sites predicted in 2 replicates of sequencing of 100% m$^6$A-modified curlcakes. **(C)** Replicability between predicted modification frequency in 2 replicates of sequencing of 100% m6A-modified curlcakes (axes are log-scaled).

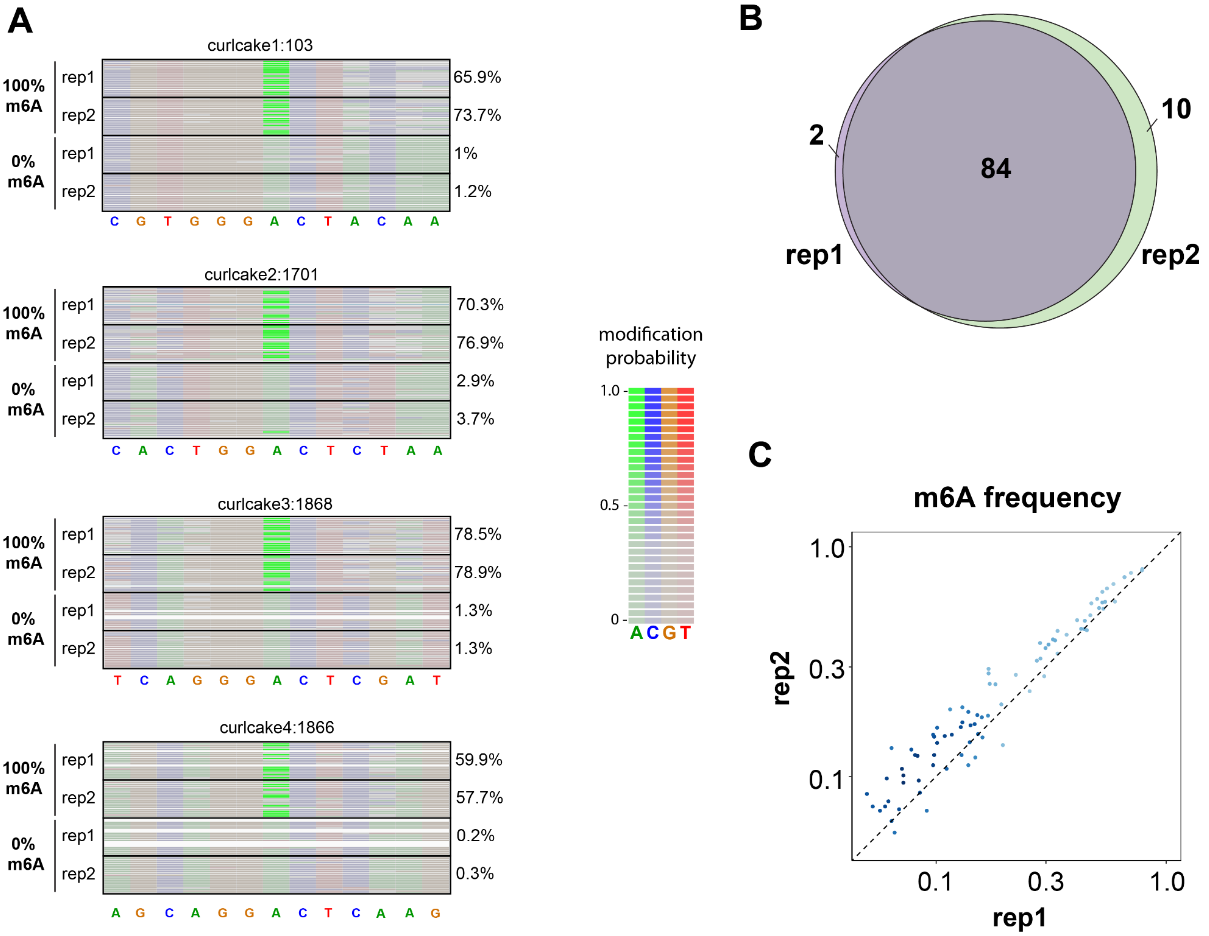

**Figure S2. Characterization of m$^6$A sites predicted by *m$^6$ABasecaller* in human (HEK293T), mouse (mESC) and zebrafish (4hpf embryos). (A,B)** Metagene plot of the distribution of m6A sites along coding transcript features, in mESC (A) and zebrafish (B) publicly available DRS datasets. **(C,D)** Venn diagram depicting the intersection between predicted m$^6$A sites in 2 independent biological replicates of HEK293T (C) and mESC (D) DRS datasets. A site was defined as "m$^6$A-modified" if it had a minimum coverage of 25 reads and a modification stoichiometry greater or equal than 5%. **(E,F)** Scatterplots depicting the correlation of modification stoichiometry of m$^6$A sites in HEK293T (E) and mESC (F) WT samples. Each dot represents an m$^6$A site. Only m$^6$A sites with more than 5% modification frequency and coverage greater or equal than 25 reads of coverage (in mESC, panel E) or 50 reads coverage (in HEK293T, panel F) in both replicates were included in the analysis. See also **Figure 3D** for scatterplot of modification frequencies across HEK293T WT replicates with 25 reads coverage threshold.

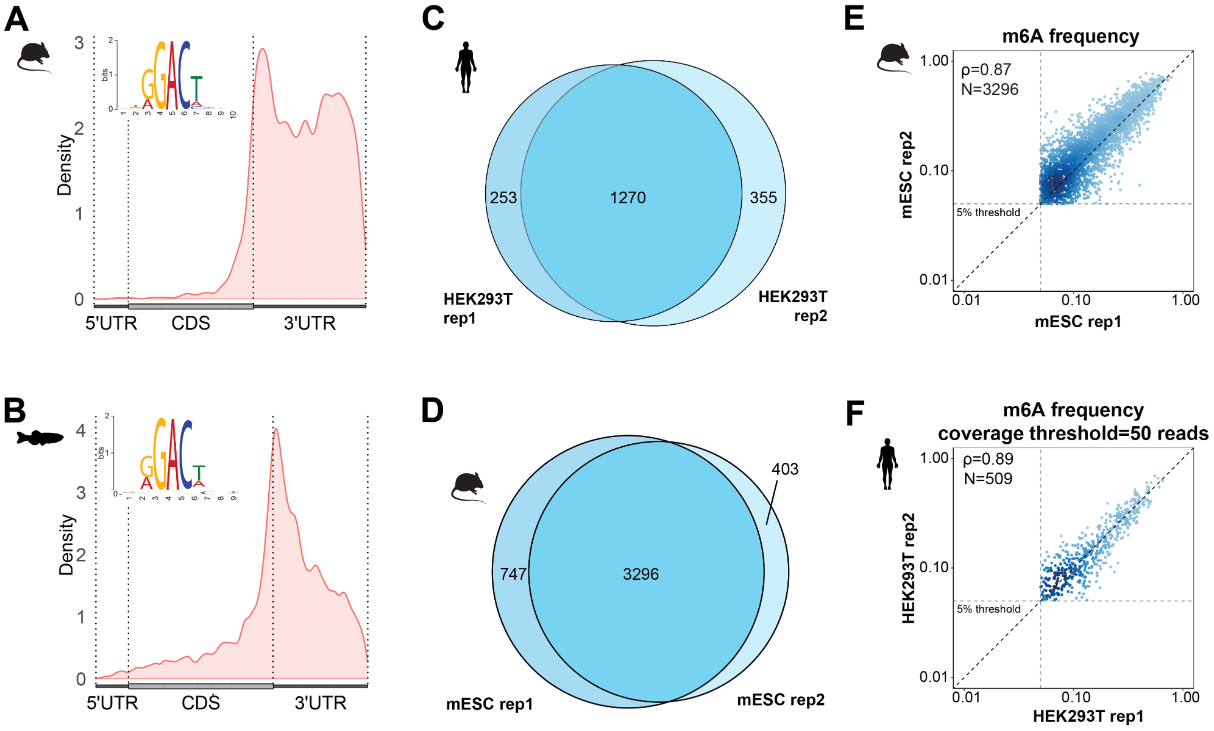

**Figure S3. m6ABasecaller captures quantitative changes in m6A stoichiometry in mES cells upon METTL3 or METTL14 KO. (A,B)** $m^6A$ modification frequencies in WT vs METTL3 KO (A) and in WT vs METTL14 KO (B) mESc DRS samples. Vertical and horizontal dashed lines denote the 5% threshold applied for a given site to be predicted as '$m^6A$-modified'. Both axes are $log_{10}$-scaled for enhanced visualization. **(C)** Density plot distribution of $m^6A$ modification frequencies in HEK293T WT, METTL3 KO and IVT samples, in two independent biological replicates**.** Dashed vertical lines represent the median $m^6A$ modification frequency observed in each sample (rep1 WT: 15.1%, rep2 WT: 15.6%, rep1 KO: 2.8%, rep2 KO: 2.9%, rep1 IVT: 0%, rep2 IVT: 0%). **(D)** Density plot distribution of m6A modification frequencies in mESC in WT, METTL3 KO and METTL14 KO samples**.** Dashed vertical lines represent the median $m^6A$ modification frequency observed in each sample (rep1 WT: 18.8%, rep2 WT: 16.4%, METTL3 KO: 4.7%, METTL14 KO: 7.8%). For C and D, a pseudocount of 0.001 was added to all values to allow logarithmic scaled axes.

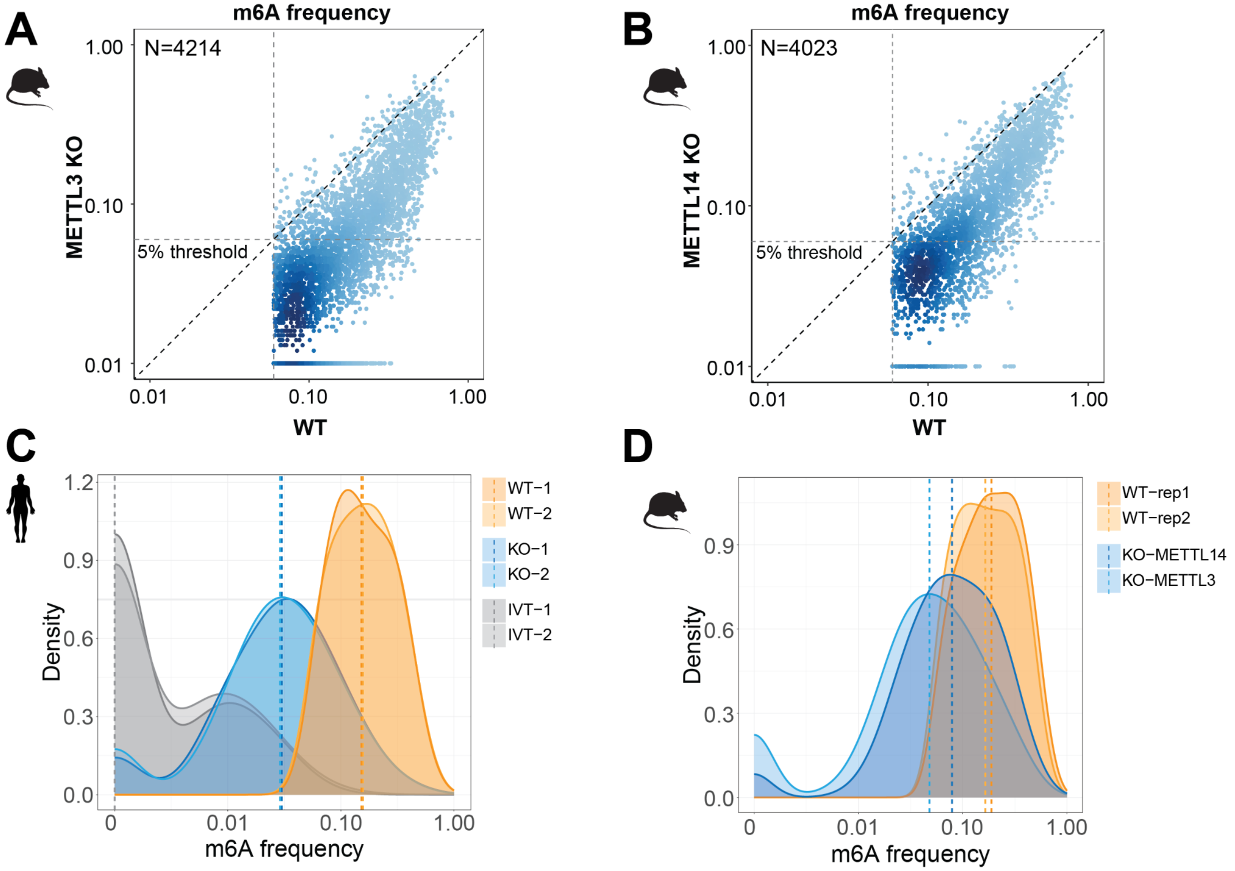

**Figure S4. Overlap between *m⁶ABasecaller* and orthogonal methods. (A,B,C)** Overlap between $m^6A$ sites found in GLORIseq and *$m^6$ABasecaller* (A) m6ACE-seq and *$m^6$ABasecaller* (B) and miCLIP and *$m^6$ABasecaller* (C) in HEK293T cells. **(D)** Overlap between predicted $m^6A$ sites using 3 different Illumina-based orthogonal methods (GLORI-seq, miCLIP and m6ACEseq) in HEK293T cells.

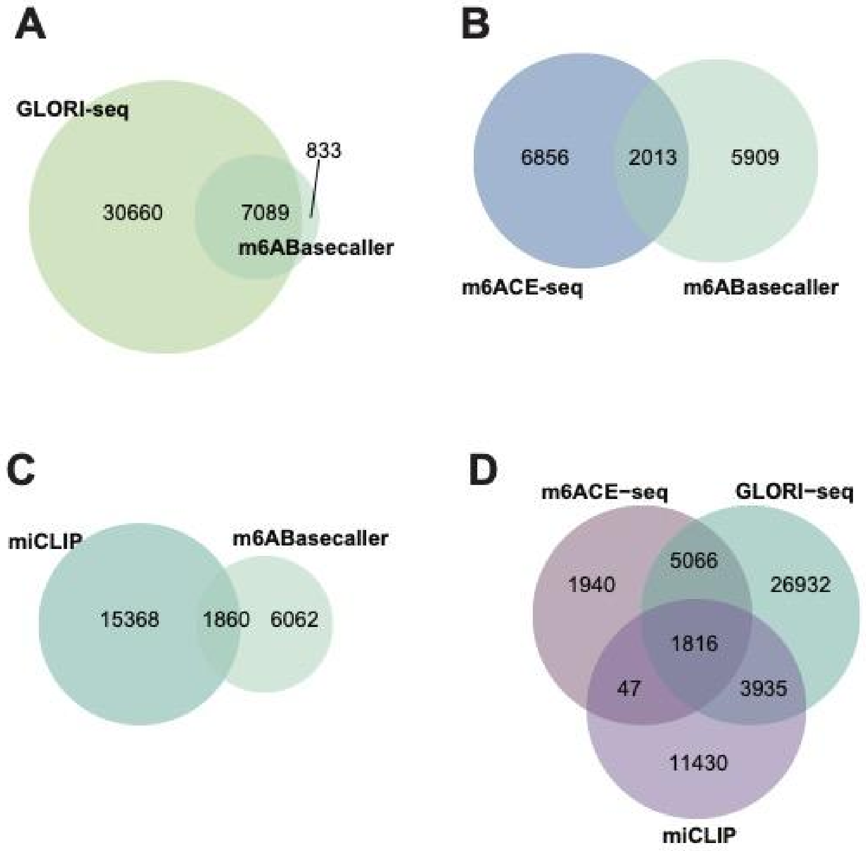

**Figure S5. Changes in m$^6$A modification stoichiometry in mESC mRNAs upon STM2457 treatment. (A)** Euclidean distance matrix calculated on the modification frequencies measured when using different concentrations of STM2457 in 2 independent replicates. **(B)** Scatterplot depicting the replicability of m$^6$A modification frequencies predicted, for untreated (CTR) and treated samples, with 3 different inhibitor concentrations: 2uM, 10uM and 20uM. Axes are log$_{10}$-scaled. **(C)** MEME motif obtained using as input the sequence context of predicted m$^6$A sites in mESC untreated (CTR) samples. **(D)** Metagene plot of the distribution along coding transcripts (N of genes=385) of the m$^6$A sites found in CTR pooled samples (N of sites=584) with >=25 reads of coverage and >=5% frequency.

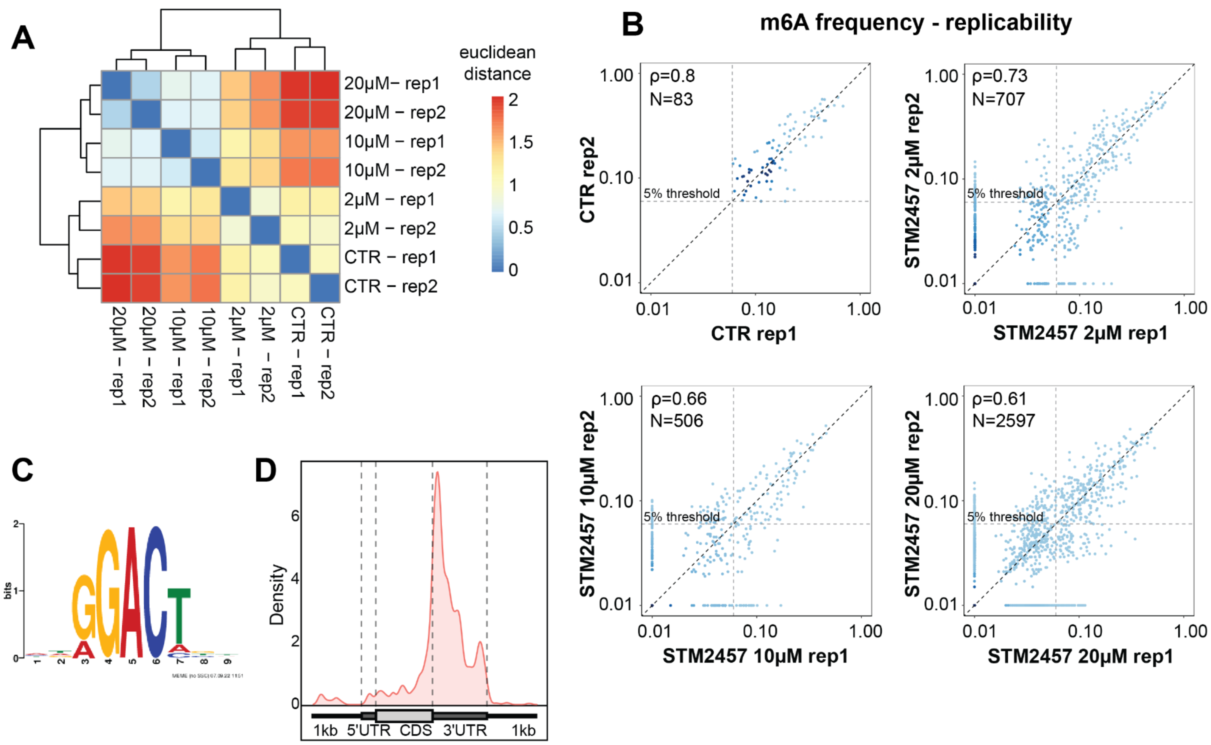

**Figure S6. Replicability of m$^6$A frequencies in tamoxifen-treated and untreated mESC cells. (A)** Replicability of m$^6$A frequencies in two replicates of mESC treated with tamoxifen (METTL3 KO) or vehicle MetOH (CTR) for 6 days. Both axes are $\log_{10}$-scaled. **(B)** Replicability of m$^6$A frequencies in the two replicates of mESC cells treated with tamoxifen (KO) or vehicle MetOH (CTR) for 14 days. Both axes are $\log_{10}$-scaled.

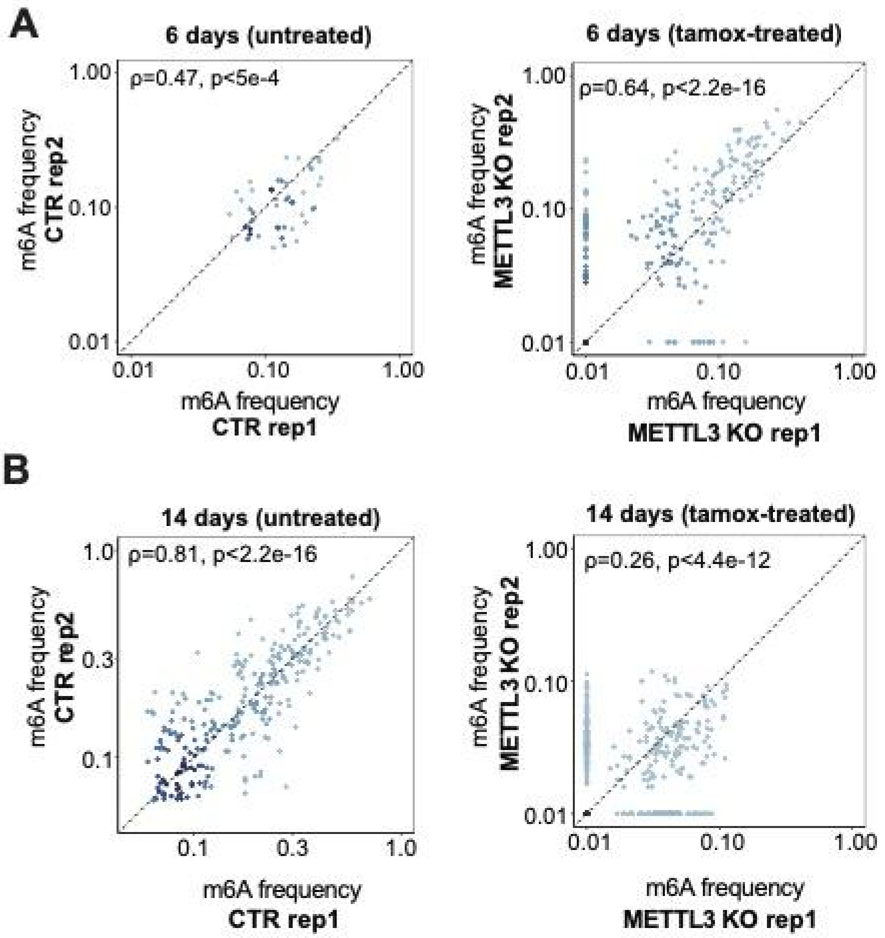

**Figure S7. Per-read analysis of m⁶A-modified sites in HepG2 cells shows dependency between m⁶A presence and polyA tail length.** Cumulative fraction of polyA tail length across reads that are binned based on the number of m6A-modified sites per read (as predicted by the *m⁶ABasecaller*). Only full-length reads are included in the analysis. The number of reads included in each bin is shown in the legend.

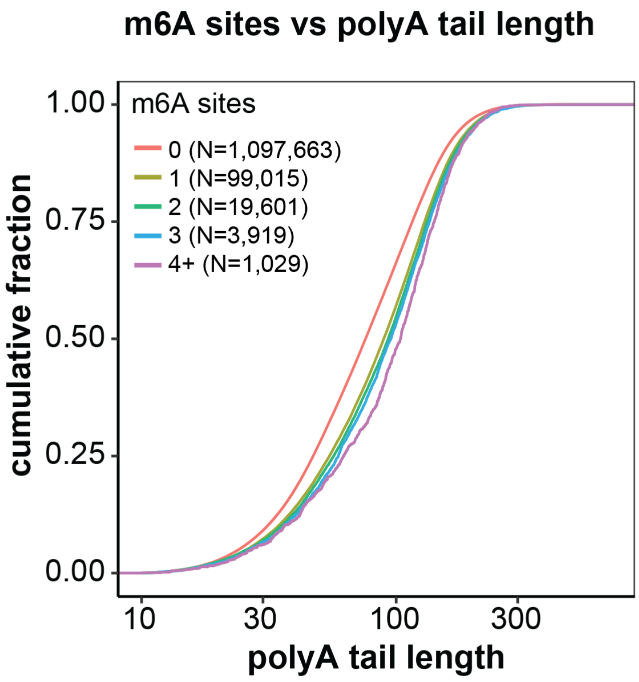

**Figure S8. Per-isoform analysis of m6A modifications. (A)** IGV snapshot of two isoforms belonging to gene UQCRFS1. In the zoomed section, the presence of m$^6$A at per read level is shown with bright red color (as the reads map to the "-" strand). Clustering of reads belonging to reassigned isoforms ENST00000304863_0 (left) or ENST00000304863_1 (right) based on their modification pattern in positions chr19:29207544 and chr19:29207538. Next to both snapshots, results from the co-occurrence analysis are shown. **(B)** IGV snapshot depicting the modification frequency at position chr9:136,862,324 in two isoforms from gene EDF1. Modification frequency at per-isoform level is shown in two replicates.

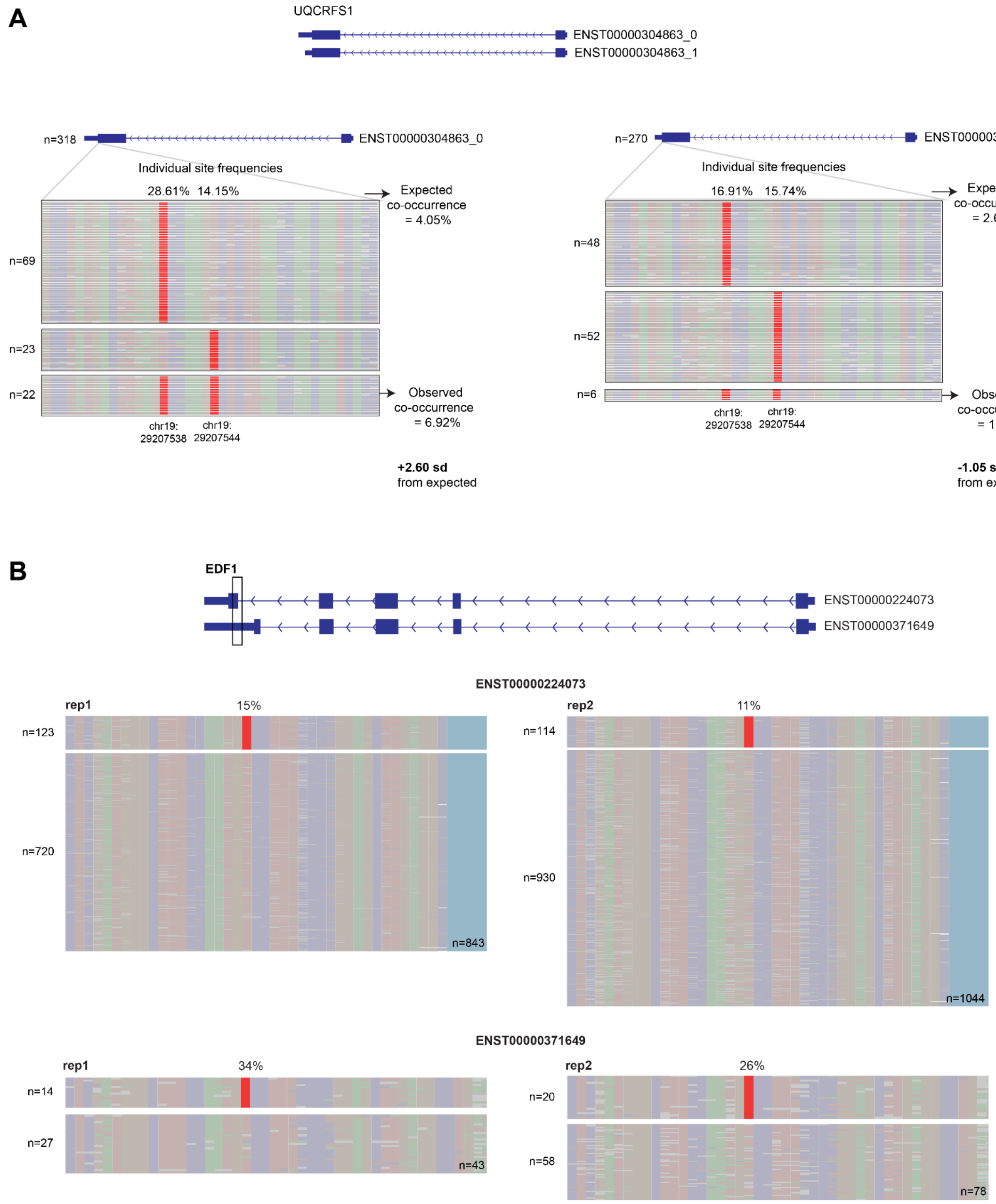
